# Exosomes exploit the virus entry machinery and pathway to transmit IFN-α-induced antiviral activity

**DOI:** 10.1101/300715

**Authors:** Zhenlan Yao, Xiaofang Li, Jieliang Chen, Yunsheng Qiao, Fang Shen, Bisheng Shi, Jia Liu, Jiahui Ding, Lu Peng, Jianhua Li, Zhenghong Yuan

## Abstract

Interferon-α (IFN-α) induces the transfer of resistance to hepatitis B virus (HBV) from liver nonparenchymal cells (LNPCs) to hepatocytes via exosomes. However, little is known about the entry machinery and pathway involved in the transmission of IFN-α-induced antiviral activity. Here, we found that macrophage exosomes depend on T cell immunoglobulin and mucin receptor 1 (TIM-1), a hepatitis A virus (HAV) receptor, to enter hepatocytes for delivering IFN-α-induced anti-HBV activity. Moreover, two primary endocytic routes for virus infection, clathrin-mediated endocytosis (CME) and macropinocytosis, collaborate to permit exosome entry and anti-HBV activity transfer. Subsequently, lysobisphosphatidic acid (LBPA), an anionic lipid closely related to endosome penetration of virus, facilitates membrane fusion of exosomes in late endosomes/ multivesicular bodies (LEs/MVBs) and the accompanying exosomal cargo uncoating. Together, this study provides comprehensive insights into the transmission route of macrophage exosomes to efficiently deliver IFN-α-induced anti-HBV activity and highlights the similarities between the entry mechanisms of exosomes and virus.

**Importance:** Our previous study showed that LNPC-derived exosomes could transmit IFN-α-induced antiviral activity to HBV replicating hepatocytes, but the concrete transmission mechanisms which include exosome entry and exosomal cargo release remain unclear. In this study, we found that virus entry machinery and pathway were also applied to exosome-mediated cell-to-cell antiviral activity transfer. Macrophage-derived exosomes exploit hepatitis A virus receptor for access to hepatocytes. Later, CME and macropinocytosis are utilized by exosomes which is followed by exosome-endosome fusion for efficient transfer of IFN-α-induced anti-HBV activity. Dissecting the similarities between exosome and virus entry will be beneficial to designing exosomes as efficient vehicles for antiviral therapy.

## Introduction

Hepatitis B virus (HBV) is a small, enveloped DNA virus that replicates via an RNA intermediate and belongs to the Hepadnaviridae family(1). Approximately 400 million people are chronically infected with HBV worldwide(2). Chronic HBV infection is a major risk factor for the development of liver cirrhosis and hepatocellular carcinoma(3). Interferon (IFN)-α is licensed for the treatment of HBV chronic infection, with a response rate of 30-40% and a clinical cure rate of approximately 10%(4), but its efficacy is limited in hepatocytes (5, 6). We and others previously reported that IFN-α induced the transfer of resistance to hepatitis viruses from nonpermissive liver nonparenchymal cells (LNPCs), including liver resident macrophages, to permissive hepatocytes via exosomes, but the underlying mechanism remains largely unclear(7–11).

Exosomes are 40–100 nm membrane vesicles derived from the intraluminal vesicles (ILVs) of multivesicular bodies (MVBs) that are secreted into the extracellular milieu through the fusion of MVBs with the plasma membrane(12, 13). These vesicles can serve as mediators of intercellular communication to exchange functional proteins, lipids, mRNAs and microRNAs (miRNAs) among cells(14–16). Given the emerging roles of exosomes from IFN-α-induced LNPCs in the antiviral innate response and their therapeutic potential(7, 8, 17, 18), it is important to understand the molecular mechanisms by which nonparenchymal cell-derived exosomes are taken up into hepatocytes and release their cargo to inhibit HBV replication.

The entry strategy used by a given exosome may depend on the proteins and lipids on the surfaces of both exosomes and recipient cells(19–21). The routes and fates of exosome internalization may partially overlap with those of the virus(9, 22, 23). Here, we found that the hepatitis A virus receptor, TIM-1, mediated the internalization of macrophage-derived exosomes into hepatocytes; we showed that the rapid clathrin-dependent pathway in concert with sustained macropinocytosis, two primary pathways for virus invasion, were also used as the major endocytic routes for exosome entry and the transmission of IFN-α-induced HBV resistance. After internalization, membrane fusion of exosomes and accompanying exosomal cargo uncoating took place in LEs/MVBs, relying on the LE-specific anionic lipid lysobisphosphatidic acid (LBPA). Collectively, our findings demonstrate that macrophage exosomes require virus entry machinery and pathway for transmission of IFN-α-induced antiviral activity to combat HBV in hepatocytes.

## Materials and methods

### Antibodies, reagents and Chemical Inhibitors

Antibodies for LAMP-2 (sc-18822), EEA1 (sc-33585) and normal mouse IgG (sc-2025) were purchased from Santa Cruz Biotechnology. Antibodies for Alix (12422-1-AP), TSG101 (14497-1-AP), CD63 (25682-1-AP), RAB5 (11947-1-AP) and RAB7 (55469-1-AP) were purchased from Proteintech Group (Rosemont, USA). Antibody for clathrin heavy chain (ab21679) was from Abcam (Cambridge, USA). Antibodies for β-actin (A2228) and GFP (G6539) were from Sigma-Aldrich. Antibody for LBPA (MABT837) was from EMD Millipore (Billerica, USA). Fluorescent secondary antibodies (A11001, A10523) were purchased from Invitrogen. Annexin V -FITC (640905) was purchased from Biolegend (San Diego, USA). Phalloidin-iFluor 488 (23115) was from AAT Bioquest (Sunnyvale, USA). Fc-TIM-1-His, a protein of TIM-1 extracellular domain (AAC39862.1) (Ser 21-Gly 290) which is fused with a polyhistidine tag at the C-terminus and the Fc region of human IgG1 at the N-terminus was from Sino Biological (Beijing, China). Aldehyde/Sulfate Latex Beads(4% w/v, 4 μm) was from Invitrogen (Carlsbad, USA).

Chemical inhibitors including dynasore (D7693), MβCD (C4555), EIPA (A3085), IPA-3 (I2285) and rottlerin (R5648) were from Sigma-Aldrich. Filipin III (70440) was purchased from Cayman chemical (Ann Arbor, USA). Chlorpromazine (S2456) and nystatin (S1934) were purchased from Selleck Chemicals (Houston, USA).

### Cells, plasmids, siRNAs and transfection

The HepG2.2.15, HepG2 and THP-1 cells used in this study have been described previously(7, 24). HepG2.2.15 and HepG2 cells were cultured in DMEM with 10% fetal bovine serum (FBS) (Biologic Industries, Beit Haemek, Israel) and Penicillin-Streptomycin (Invitrogen, Carlsbad, USA), while THP-1 cells were maintained in RPMI-1640 with 10% FBS and antibiotics. To obtain macrophage-like cells that closely resembled human monocyte-derived macrophages, THP-1 cells were differentiated via PMA stimulation (phorbol 12-myristate 13-acetate; Sigma-Aldrich, Taufkirchen, Germany), as described previously(7, 25).

Markers of endosomal compartments fused with cyan fluorescent protein (CFP), including CFP-RAB5, CFP-RAB7 and CFP-CD63, were kindly provided by Walther Mothes from Yale University in New Haven, CT, USA(26). K44A dynamin-2 pEGFP was a gift from Sandra Schmid (Addgene plasmid # 34687). pcDNA3-EGFP-Cdc42-T17N (Addgene plasmid # 12976) and pcDNA3-EGFP-Rac1-T17N (Addgene plasmid # 12982) were gifts from Gary Bokoch. Caveolin-1 labeled with C-terminal tag of enhanced green fluorescent protein (EGFP) was constructed by insertion of the claveolin-1 cDNA fragment into a pEGFP-N1 expression vector (Clontech, Palo Alto, USA). To produce GFP-carrying exosomes, a THP-1 cell line stably expressing GFP was established via lentivirus transduction. The lentiviral vector PLJM1-GFP (Addgene) was used to generate lentivirus for the transduction, according to the manufacturer’s instructions (Addgene). Stable GFP-expressing THP-1 cells were selected by flow cytometric sorting (BD FACSAria II; BD Biosciences, San Jose, USA). siRNAs for clathrin heavy chain, caveolin-1 and negative control were purchased from Santa Cruz Biotechnology. siRNA for TIM-1 was purchased from Ruibo.

DNA plasmid transfection into HepG2 cells was performed using Lipofectamine 2000 (Invitrogen). For RNA-mediated interference, HepG2 cells at 30 to 40% confluence were transfected with 50 nM small interfering RNA (siRNA) duplexes designed and purchased from Santa Cruz (Dallas, USA) or Ruibo (Guangzhou, China) using RNAiMAX (Invitrogen) according to the manufacturer’s instructions. At 24 h post-transfection, the cells were transfected again with 50 nM of the same siRNA duplexes. The following treatment was performed 72 h after the first siRNA transfection.

### Exosome purification, characterization and labeling

Macrophages derived from THP-1 or GFP-expressing THP-1 cells were grown in culture medium supplemented with 10% FBS (which was depleted of endogenous exosomes by overnight centrifugation at 100,000 g). Exosomes from the culture supernatants were isolated by differential centrifugation, as described previously(7). To obtain exosomes from IFN-α-treated macrophages, the macrophages were treated for 48 h with 1,000 U/ml of IFN-α (PBL Assay Science, New Brunswick, USA) before isolation. The purified exosomes were characterized via electron microscopy and immunoblot analysis, as described previously(7). Protein amounts of exosomes were quantified using a BCA protein assay kit (Pierce, Rockford, USA).

The isolated exosomes were labeled with PKH67 or PKH26 according to the manufacturer’s protocol (Sigma-Aldrich) for use in endocytosis assays. For the fluorescence self-quenching assay for membrane fusion, R18 (Octadecyl Rhodamine B Chloride, Invitrogen) was inserted into the viral membranes at a self-quenching surface density(27, 28).

### Endocytosis assays of exosomes

To assay exosome internalization, 10-20 μg/ml of labeled exosomes were added to HepG2 cells cultured with serum-free medium and incubated at 37°C. HepG2 cells were untreated or pretreated with the indicated amounts of inhibitors for 30 min before incubation with exosomes or endocytic markers. Except cholesterol inhibitors (MβCD, nystatin, and filipin III), inhibitory compounds were present continuously during subsequent endocytosis assays. Despite moderate cytotoxicity of MβCD-treated cells, no significant toxicity was observed for the other inhibitors (data not shown), which indicated that inhibition of exosome internalization was not caused by cytotoxicity. As controls, HepG2 cells were incubated with 2 μg/ml of Alexa568-transferrin (Invitrogen) or 0.2 mg/ml of dextran labeled with Rhodamine B isothiocyanate (RhoB-dextran, Sigma-Aldrich) for 30 min or 1 h at 37°C. For competitive inhibition of TIM-1-mediated exosome entry by Fc-TIM-1-His, HepG2 cells were incubated with labeled exosomes in presence of 1 μg/ml Fc-TIM-1-His at 37°C for 2 h. Endocytosis was stopped, and surface-bound exosomes or markers were removed by washing with ice-cold PBS.

### Confocal laser-scanning and time-lapse microscopy

Confocal images were captured using a Leica TCS SP8 confocal microscope (Leica Microsystems, Buffalo Grove, USA) with a 400X or 630X oil objective (pinhole set at 1 Airy unit) and processed using LAS X (Leica). For time-lapse microscopy analysis, HepG2 cells were grown in 35-mm glass bottom culture dishes with four chambers (Cellvis, Mountain View, USA) overnight. Before microscopic examination, the medium was changed to serum-free DMEM, and fluorescence-labeled exosomes were added and kept in the medium during image collection. Time-lapse images were captured every 10 min for 6-μm slices using a DeltaVision Elite high-resolution microscope (Applied Precision, Issaquah, USA) connected to a 37°C incubator and buffered with 5% CO_2_. The images were further processed with softWoRx Explorer (Applied Precision, Issaquah, USA) and analyzed with ImageJ (NIH, USA). For colocalization studies, the distribution patterns of the fluorescent signals were analyzed using the Plot Profile analysis tool of ImageJ, and Pearson’s correlation coefficients (Rr) were obtained by using the Colocalization finder plugin of ImageJ. For the Pearson’s correlation coefficients (Rr), the values ranged from 1 (a perfect positive correlation) to –1 (a perfect negative correlation), with 0 representing a random distribution(29). Time-related
fluorescence intensities of the R18 dequenching signals were assessed using the Time Series Analyzer V3 plugin of ImageJ.

### Flow cytometry analysis

For endocytosis assay, cells were washed three times with ice-cold PBS, detached using trypsin, and subsequently resuspended in PBS with 1% FBS. Flow cytometry analysis was performed on an LSR Fortessa instrument integrated with the FACSDiva software (BD Biosciences). A minimum of 10,000 events within the gated live cells were collected and analyzed per sample using FlowJo (Tree Star, Ashland, USA).

For PtdSer detection, 4 μm latex beads were coated with exosomes through 2-hour incubation at room temperature. The exosome-bead complexes were then blocked with 200 mM glycine and normal IgG and washed with 1% FBS which was followed by annexin V-FITC labeling for 40 min at 4°C. The exosome-bead complexes were subsequently washed and suspended with 1% FBS for flow cytometry analysis. A minimum of 50,000 events within the gated exosome-bead complexes were collected and analyzed per sample via FlowJo.

### HBV DNA quantitation and antigen measurement

HepG2.2.15 cells pre-transfected with siRNAs or pretreated with chemical inhibitors were incubated with exosomes isolated from macrophages with or without IFN-α treatment at a concentration of 10 μg/ml for 48 h. The supernatant of the HepG2.2.15 culture was collected and transferred for viral antigen measurement using the enzyme-linked immunosorbent assay (Kehua ELISA kit; Kehua, Shanghai, China). HBV DNA levels in the culture medium were extracted using a MagNA Pure 96 system (Roche, Shanghai, China) and quantified using real-time PCR.

### Statistics

All data are presented as the mean of duplicates ± S.D. Statistical comparisons were made using a two-tailed Student’s t-test; *P* values of 0.05 or less were considered to be statistically significant.

## Results

### PtdSer receptor TIM-1 is necessary for exosome entry and the transfer of IFN-α-induced anti-HBV activity

Exosomes were isolated from the culture of THP-1-derived macrophages by differential centrifugation, as described previously(7). Membrane vesicles approximately 100 nm in diameter with a cup-shaped structure typical of exosomes were identified by electron microscopy (Fig. 1A). Further characterization by immunoblotting indicated the presence of exosomal markers (CD63, TSG101, and Alix), conserved exosomal proteins (LAMP-2, β-actin) and the absence of the endosomal marker EEA1 (Fig. 1B). Isolated exosomes were labeled with the fluorescent lipid dye PKH26 or PKH67. We observed the internalization of PKH26-labeled exosomes by hepatocyte-derived HepG2 cells, which were stained for cytoskeletal F-actin with Phalloidin-iFluor 488 (Fig. 1C) at 37°C, and found that the uptake kinetics were time- and concentration-dependent (Fig. 1D).

**Figure 1.**
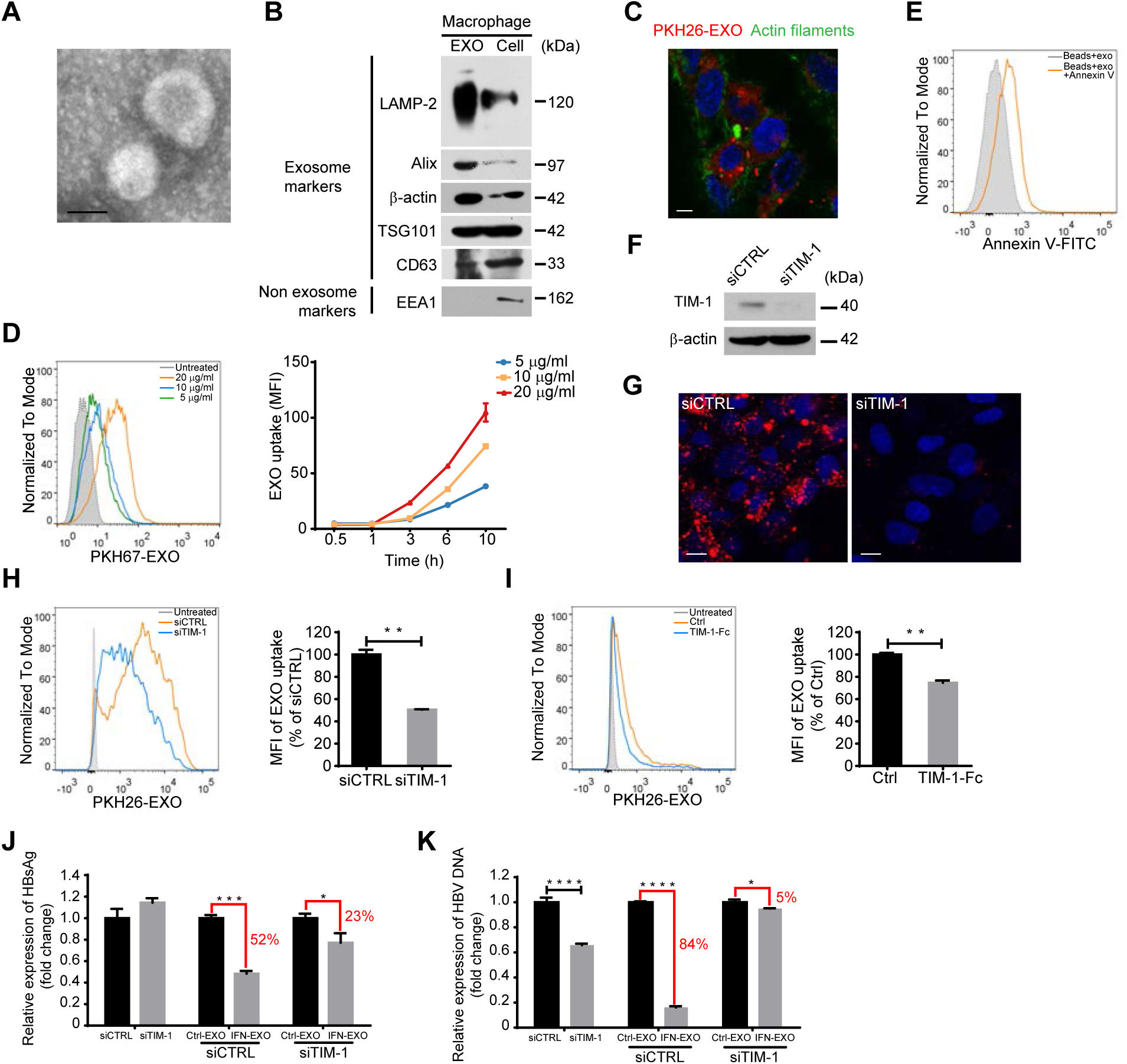
TIM-1 mediates exosome internalization and IFN-α-induced anti-HBV activity transmission. **(A)** Electron microscopy of purified exosomes from macrophages. Scale bar: 100 nm. **(B)** Immunoblot analysis of macrophage-derived exosomes (left) and corresponding cells (right) for exosomal and non exosomal markers. **(C)** PKH26-labeled exosome internalization by HepG2 cells. Scale bar: 5 μm. **(D)** Time- and concentration-dependent uptake of exosomes. HepG2 cells were incubated with PKH67-labeled exosomes (PKH67-EXO) at the indicated concentrations for up to 10 h (right). The fluorescence intensity distribution of cells incubated with PKH67-EXO for 3 h is also shown (left). **(E)** PtdSer detection on the exosome surface. Exosomes coated onto 4 μM latex beads were either stained or not with Annexin V-FITC and analyzed by flow cytometry. **(F)** Knockdown validation of TIM-1 by immunoblot. **(G, H)** Confocal images (G) or flow cytometry analysis (H) of PKH26-labeled exosome internalization by HepG2 cells after TIM-1 knockdown. Scale bars: 10 μm. For flow cytometry analysis, both histogram graph (left) and mean fluorescence intensities (MFI) (right) which are normalized to siCTRL-transfected cells are presented. **(I)** Flow cytometry analysis of PKH26-labeled exosome internalization by HepG2 cells in presence or absence (Ctrl) of Fc-TIM-1-His. MFI (right) are normalized to ctrl cells. **(J, K)** Blockade of IFN-α-induced anti-HBV activity transmission by TIM-1 knockdown. HepG2.2.15 cells transfected with either siTIM-1 or siCTRL were treated with exosomes from IFN-α-stimulated macrophages (IFN-EXO) or unstimulated cells (Ctrl-EXO). HBsAg and HBV DNA levels in the medium were measured by ELISA (J) or quantified by qPCR (K). The error bars indicate the SD. ^⋆^*P* < 0.05, ^⋆⋆^*P* < 0.01, ^⋆⋆⋆^*P* < 0.001, ^⋆⋆⋆⋆^*P* < 0.0001 (Student’s t-test). The data are representative of three independent experiments.

PtdSer — an apoptosis marker typically located on the inner leaflet of the plasma membrane — is found on the outer membrane of exosomes from bone marrow derived dendritic cells (BMDCs) and oligodendrocytes(20, 30). Previous experimental evidence indicates that some viruses may exploit PtdSer as apoptotic disguise and enter target cells through PtdSer receptor-mediated internalization(31). To determine whether and which PtdSer receptors play a role in the entry of macrophage-derived exosomes into hepatocytes, we first confirmed PtdSer expression on the outer membrane of macrophage-derived exosomes through annexin-V labeling of exosomes isolated from macrophages (Fig. 1 E).

We then inhibited the expression of two hepatic PtdSer receptors involved in virus entry(31), T cell immunoglobulin and mucin receptor 1 (TIM-1) (Fig. 1F) and Growth Arrest Specific 6 (GAS6) (data not shown), in HepG2 cells with specific siRNAs. The uptake of PKH26-labeled exosomes was significantly reduced in HepG2 cells after TIM-1 knockdown (Fig. 1G and H), but interference via GAS6 expression had no effect on exosome uptake (data not shown). It is notable that the IgV in ectodomains of TIM proteins bind PtdSer on viral envelope and enhance virus entry(32). Exogenous Fc-TIM-1-His, TIM-1 extracellular domain fused with His tag and Fc region of human IgG1, competitively inhibited exosome internalization by HepG2 cells (Figure 1I), which suggested that the ectodomain of TIM-1 also play a functional role in exosome entry. Corresponding to previous results reflecting the engagement of TIM-1 in exosome uptake, IFN-α-induced anti-HBV activity mediated by exosomes from IFN-α-stimulated macrophages (IFN-EXO) was diminished in TIM-1-knockdown HepG2.215 cells in comparison to that in cells transfected with control (ctrl) siRNA, as indicated by HBsAg expression (Fig. 1J). In addition, IFN-α-induced exosome-mediated antiviral activity only slightly suppressed HBV DNA production in the supernatant of TIM-1-knockdown cells, in contrast to cells transfected with ctrl siRNA (Fig. 1K). It was unexpected that the knockdown of TIM-1 caused a decrease in HBV DNA in the supernatant, which suggests that TIM-1 is a positive factor for HBV replication (Fig. 1K). Collectively, these findings demonstrated that PtdSer and its receptor TIM-1 act as portals for exosomal internalization and the transfer of IFN-α-induced antiviral activity against HBV.

### Dynamin-2 and cholesterol are required for exosome entry into hepatocytes

The interaction of exosomes with receptors on donor cells can induce the cellular response of internalization through endocytic pathways(33). Endocytosis occurs via several pinocytic mechanisms that include the clathrin-mediated mechanism, macropinocytosis, the caveolae-mediated mechanism and other less well-defined mechanisms(34, 35). The large GTPase dynamin-2 functions at the heart of endocytic vesicle fission in clathrin-mediated endocytosis (CME) and caveolae-mediated endocytosis (Fig. 2A)(36). Recent studies showed that dynamin is also responsible for the closure of circular ruffles in macropinocytosis (Fig. 2A)(37). Cholesterol plays essential roles in the formation of caveolae, clathrin-coated pit budding and membrane ruffling in macropinocytosis (Fig. 2A)(38–40).

**Figure 2.**
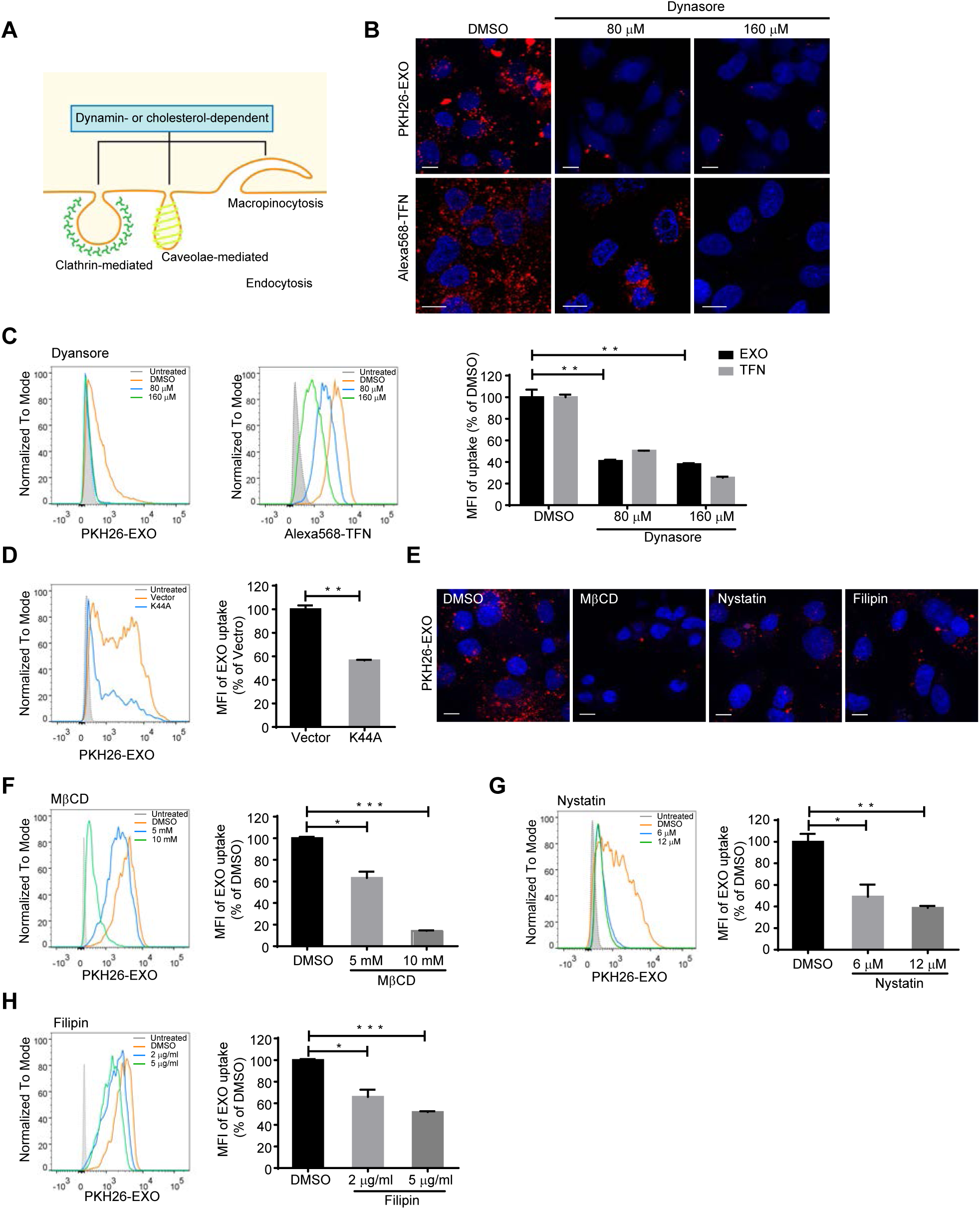
Exosome internalization is dynamin- and cholesterol-dependent. **(A)** Schematic representation of the roles of dynamin-2 and cholesterol in various endocytic pathways. **(B, C)** Confocal images (B) or flow cytometry analysis (C) of exosome and transferrin internalization by HepG2 cells treated with dynasore. Scale bars: 10 μm. MFI (right) are normalized to DMSO-treated cells. **(D)** Flow cytometry analysis of exosome internalization by HepG2 cells transfected with EGFP-Dyn2K44A mutant. HepG2 cells transfected with EGFP-tagged dominant-negative dynK44A mutant were incubated with PKH26-labeled exosomes. Transfected cells (EGFP+) are gated, and the uptake of exosomes among transfected cells (EGFP+/PKH26+) is analyzed and presented by histogram graph (left) and MFI (right). MFI are normalized to vector-transfected controls. **(E-H)** Confocal images (E) or flow cytometry analysis (F-H) of exosome internalization by HepG2 cells treated with cholesterol inhibitors (MβCD, Nystatin and Filipin). Scale bars: 10 μm. For flow cytometry analysis, MFI (right) are normalized to DMSO-treated cells. The error bars indicate the SD. ^⋆^*P* < 0.05, ^⋆⋆^*P* < 0.01, ^⋆⋆⋆^*P* < 0.001 (Student’s t-test). The data are representative of three independent experiments.

To investigate the role of dynamin-2 in exosome entry, we suppressed the function of dynamin-2 in HepG2 cells with the specific inhibitor dynasore. The efficacy of dynasore was confirmed using Alexa568-labeled transferrin (Alexa568-TFN), which is the best-characterized cargo protein of CME (Fig. 2B and C). The uptake of PKH26-labeled exosomes was reduced by approximately 60% following dynasore treatment (Fig. 2C). In addition, the expression of the dominant-negative mutant of dynamin-2, Dyn2K44A, also significantly blocked exosome entry (Fig. 2D). We next sought to determine whether cholesterol is necessary for exosome entry into hepatocytes. Using Methyl-β-cyclodextran (MβCD) to extract cholesterol from the plasma membrane of HepG2 cells significantly inhibited PKH26-labeled exosome entry (Fig. 2E and F). The reduction was up to 86% when treating HepG2 cells with 10 mM MβCD (Fig. 2 F). Masking cholesterol with binding compounds (nystatin and filipin) resulted in milder but still apparent inhibition of exosome uptake by HepG2 cells (Fig. 2E, G and H). These results indicated that the dynamin-2- and cholesterol-dependent endocytic pathways are required for the entry of exosomes into hepatocytes.

### Clathrin-but not caveolae-mediated endocytosis is important for exosome uptake and the transmission of IFN-α-induced anti-HBV activity

CME, which is the uptake of material into cells from the surface using clathrin-coated vesicles, is the preferred route by which some PtdSer-exposing viruses enter target cells(31). To investigate the dependence of exosome entry on CME, hepatocytes were treated with chlorpromazine (CPZ), an inhibitor of clathrin-coated pit assembly. PKH26-labeled exosome uptake decreased by 34%, and as a positive control, transferrin uptake was inhibited under the same conditions (Fig. 3A and B). Moreover, knockdown of the clathrin heavy chain (CHC) also reduced exosome entry into hepatocytes by 34% (Fig. 3C and D). To further investigate the endocytic pattern engaged in exosome entry, exosomes were stained with PKH67 and administered to HepG2 cells in the presence of Alexa568-TFN. Partial colocalization of exosomes and transferrin was observed 30 min post-internalization, while little colocalization was captured 1 h after internalization, indicating rapid clathrin-dependent endocytosis during the early stage of exosome internalization (Fig. 3E and F). Scatterplots, Pearson’s correlation coefficient (Rr) and an intensity profile were used to quantify the degree of colocalization between PKH67-labeled exosomes and Alexa568-TFN. Partial colocalization between exosomes and transferrin was evidenced by scatterplots, a fraction of which were close to diagonal, and the corresponding Rr was 0.1292 (see Materials and Methods) (Fig. 3E). There were several peak superpositions in the intensity profile (Fig. 3E). Correspondingly, the downregulation of CHC expression in HBV-replicating hepatocytes weakened the IFN-α-induced anti-HBV activity transmitted by exosomes in HepG2.2.15 cells, as indicated by viral antigen expression and DNA quantification (Fig. 3G and H). In addition, caveolae-mediated endocytosis did not appear to be required for exosome internalization by hepatocytes, as indicated by the inhibition of caveolin-1 (CAV1) expression (Fig. 3I and J). Together, these data showed that clathrin-but not caveolae-mediated endocytosis contributed to exosome uptake and the transfer of IFN-α-induced HBV resistance.

**Figure 3.**
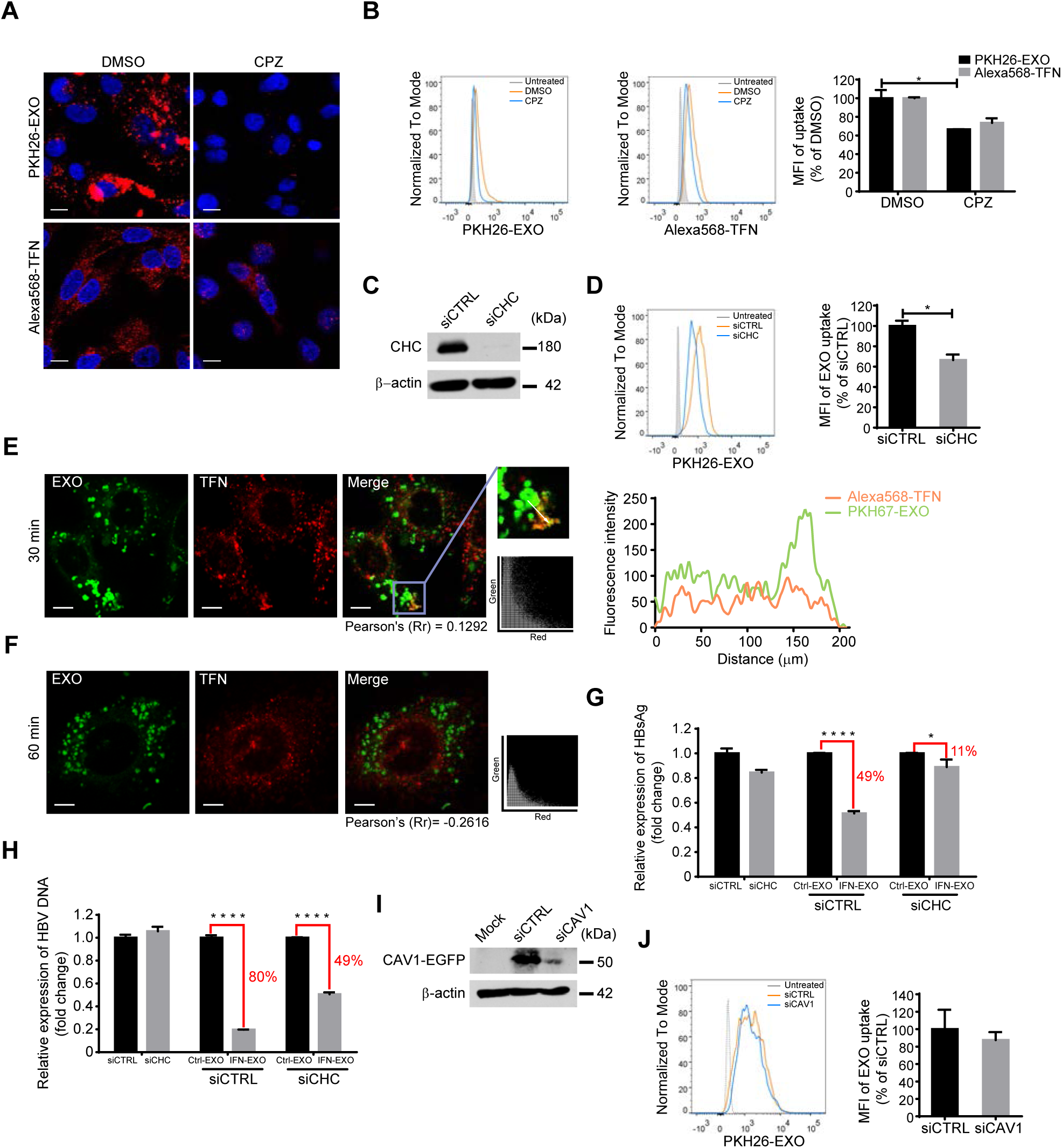
Exosome internalization involves clathrin-mediated endocytosis (CME) not caveolae-mediated endocytosis. **(A, B)** Confocal images (A) or flow cytometry analysis (B) of exosome and transferrin internalization by HepG2 cells treated with 10 μg/ml CPZ. Scale bars: 10 μm. For flow cytometry analysis, MFI (right) are normalized to DMSO-treated cells. **(C)** Knockdown validation of clathrin heavy chain (CHC) by immunoblot. **(D)** Flow cytometry analysis of exosome internalization by HepG2 cells after CHC knockdown. MFI (right) are normalized to siCTRL-transfected cells. **(E, F)** Internalized exosome colocalized with transferrin 30 (E) min or 1 h (F) after internalization. The cells were fixed and analyzed by confocal microscopy. Scatterplots and Pearson’s correlation coefficients for the overlap of red (Alexa568-transferrins) and green (PKH67-labeled exosomes) pixel intensities corresponding to the images are presented. Intensity profiles are used to describe the distribution along the indicated white arrow in the region of interest (ROI). Scale bars: 5 μm. **(G, H)** Blockade of IFN-α-induced anti-HBV activity transmission by CHC knockdown. HepG2.2.15 cells transfected with either siCHC or siCTRL were treated with IFN-EXO or Ctrl-EXO. HBsAg and HBV DNA levels in the medium were measured by ELISA (G) or quantified by qPCR (H). **I** Knockdown validation of caveolin-1 (CAV1) by immunoblot. Endogenous amount of caveolin-1 in HepG2 cells is low. To test the knock-down efficiency, siRNAs with a plasmid encoding EGFP-CAV1 were cotransfected. Expression of EGFP-CAV1 was assessed by immunoblot. **J** Flow cytometry analysis of exosome internalization by HepG2 cells with CAV1 knocked down. MFI (right) are normalized to siCTRL-transfected cells. The error bars indicate the SD. ^⋆^*P* < 0.05, ^⋆⋆⋆⋆^*p* < 0.0001 (Student’s t-test). The data are representative of three independent experiments.

### Macropinocytosis plays an alternative role in exosome uptake and the transfer of IFN-α-induced anti-HBV activity

More than one endocytic route was reported to be used in virus or exosome entry(33, 41). Given the incomplete inhibition of exosome entry by blockade of CME and the sustained increase of internalized exosomes for hours (Fig. 1D and 3B, D), there might be alternative pathways to support exosome entry into hepatocytes. Macropinocytosis is a fluid-phase type of endocytosis that is accompanied by membrane ruffles regulated by actin rearrangement(37). This process is engaged in apoptotic cell removal and is favored by some viruses that use apoptotic mimicry to enter target cells(31).

The induction of a robust increase in fluid-phase uptake is a hallmark of macropinocytosis(39). The results showed that the uptake of 70-kDa dextran labeled with Rhodamine B isothiocyanate (RhoB-dextran), which is a fluid-phase marker specific for macropinocytosis, was enhanced by incubation with macrophage-derived exosomes in HepG2 cells (Fig. 4A). A Na+/H+ exchanger (NHE) is needed for macropinosome formation via the modulation of Rho GTPases at the plasma membrane, and NHE inhibition by 5-(N-Ethyl-N-isopropyl) amiloride (EIPA) has been widely used as a diagnostic criterion for macropinocytosis(42). The entry of both exosomes and dextran into HepG2 cells was apparently inhibited by EIPA, and a remarkable decrease (80%) in exosome uptake was achieved in the presence of 80 nM EIPA (Fig. 4B and C). PAK1 and PKC are two serine/threonine kinases that are required for macropinocytosis(39). We found that exosome entry was markedly blocked by the PAK1 inhibitor IPA-3 and the PKC inhibitor rottlerin (Fig. 4D-F). PKC inhibition resulted in a more significant reduction in exosome internalization by up to 66% in hepatocytes (Fig. 4F). As a positive control, dextran internalization was greatly inhibited by the two kinase inhibitors (Fig. 4 D-F). However, the expression of a dominant-negative mutant of Rac1 or Cdc42, two common GTPases that modulate membrane ruffles, had no effect on exosome internalization (Fig. 4G and H), which suggested that macrophage exosomes might enter hepatocytes via a Rac1- or Cdc42-independent route. Next, we reinvestigated the role of macropinocytosis in exosome entry by comparing the distribution patterns of dextran and exosomes after internalization. In contrast to that seen for rapid CME-dependent exosome uptake, confocal images showed consistent colocalization of PKH67-labeled exosomes with RhoB-dextran-filled intracellular vacuoles (Fig. 4I and J). A highly overlapped distribution was observed 1 h post-exosome internalization and was confirmed by the corresponding scatterplots, colocalization coefficient and intensity profile (Fig. 4J). Furthermore, the inhibition of macropinocytosis in HepG2.2.15 cells by EIPA partially blocked the IFN-α-induced anti-HBV activity mediated by exosomes derived from IFN-treated macrophages, as indicated by viral DNA quantification (Fig. 4K). Thus, we concluded that macropinocytosis served as a sustained alternative route that was active from the early stage of exosome internalization and cooperated with CME to ensure hepatocytes the access to exosome-mediated HBV resistance.

**Figure 4.**
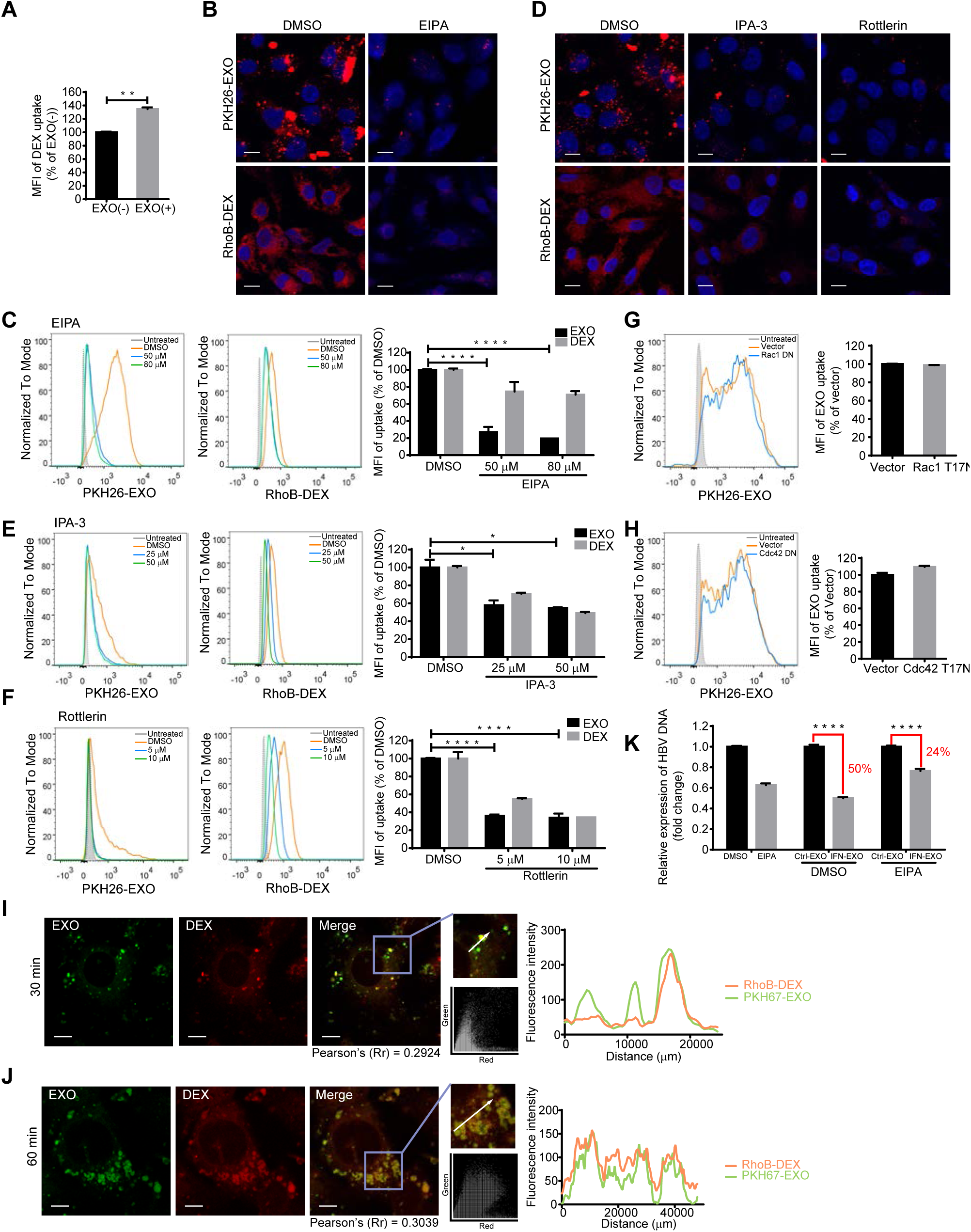
Exosome internalization involves macropinocytosis. **(A)** Preincubation with exosomes increased dextran uptake in HepG2 cells. RhoB-dextran (RhoB-DEX) uptake by HepG2 cells pretreated with exosomes (EXO(+)) was analyzed by flow cytometry, and the MFI is normalized to untreated cells (EXO(-)). **(B, C)** Confocal images (B) or flow cytometry analysis (C) of exosome and dextran internalization by HepG2 cells treated with EIPA. Scale bars: 10 μm. For flow cytometry analysis, MFI (right) are normalized to DMSO-treated cells. **(D-F)** Confocal images (D) or flow cytometry analysis (E, F) of exosome and dextran internalization by HepG2 cells treated with IPA-3 or rottlerin. Scale bars: 10 μm. For flow cytometry analysis, MFI (right) are normalized to DMSO-treated cells. **(G, H)** Exosome uptake is independent of Rac1 or Cdc42. Flow cytometry analysis of exosome internalization by HepG2 cells transfected with EGFP-Rac1 DN mutant (G) or EGFP-Cdc42 DN mutant (H), followed by the incubation with PKH26-labeled exosomes. Transfected cells (EGFP+) are gated, and the uptake of exosomes among transfected cells (EGFP+/PKH26+) is analyzed as described above. **(I, J)** Internalized exosome colocalized with dextran 30 min (I) or 1 h (J) after internalization. The colocalization of Rho-dextran (red) with PKH67-labeled exosomes (green) is analyzed as described above. Scale bars: 5 μm. **(K)** Blockade of IFN-α-induced anti-HBV activity transmission by EIPA treatment. HepG2.2.15 cells were pretreated with DMSO or EIPA which presented continuously during following incubation with IFN-EXO or Ctrl-EXO. HBV DNA levels in the medium were quantified by qPCR. The error bars indicate the SD. ^⋆^*P* < 0.05, ^⋆⋆^*P* < 0.01, ^⋆⋆⋆⋆^*P* < 0.0001 (Student’s t-test). The data are representative of three independent experiments.

### Exosomes expose cargo through membrane fusion in late endosomes/ multivesuclar bodies

Once internalized within primary endocytic vesicles, the incoming substances traffic into the endosomal system(41). The endocytosed substances are routed from early endosomes (EEs) to late endosomes (LEs, often taking the form of MVBs) and lysosomes for degradation(41). Membrane fusion-induced endosome penetration is commonly manipulated by viruses or delivery vectors to send viral genomes or biologics to the cytosol before lysosomal degradation(27, 43–45). It remains unknown whether a similar membrane fusion strategy is adopted for exosomal cargo release in endosomes after internalization (Fig. 5A).

**Figure 5.**
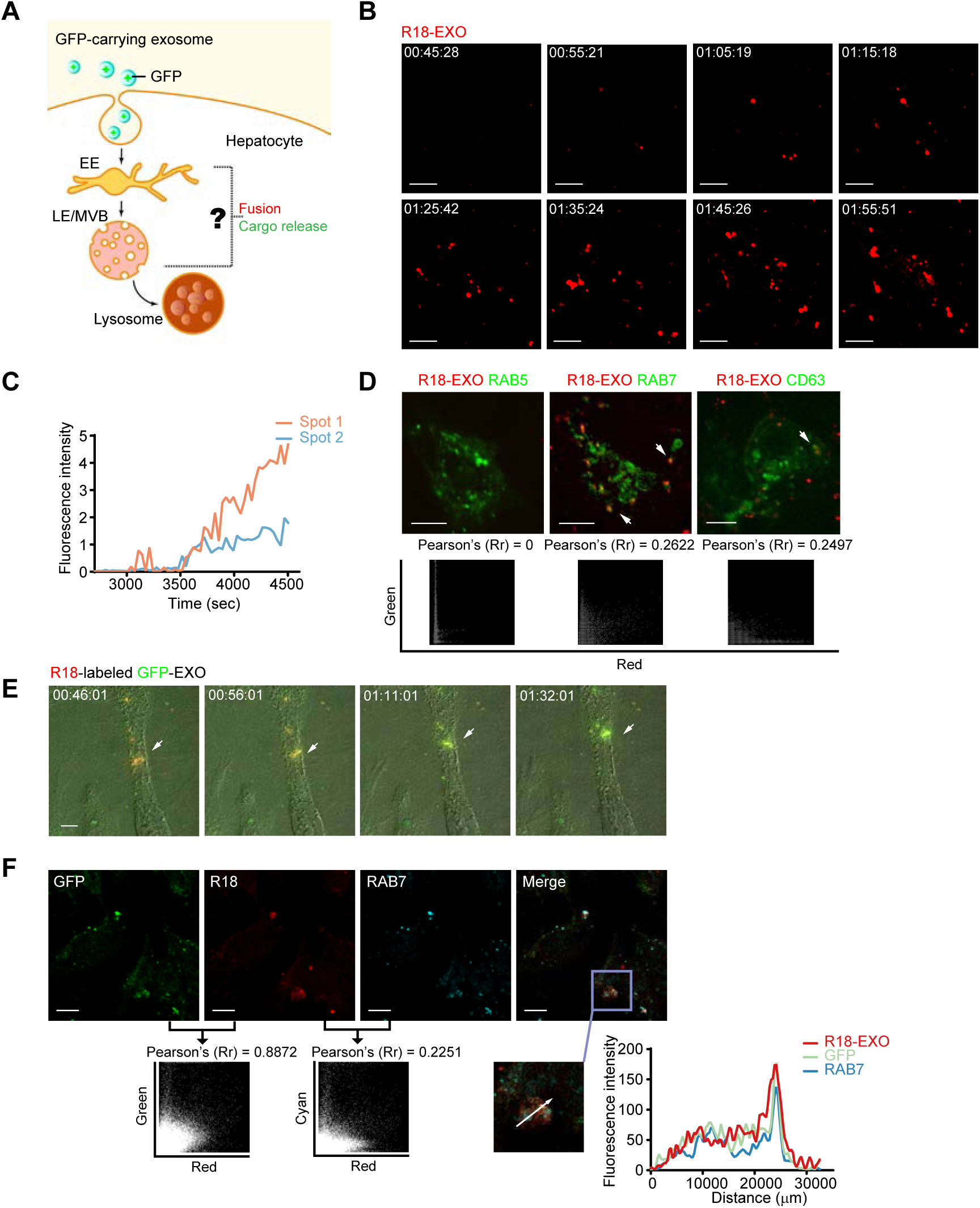
Membrane fusion of GFP-carrying exosomes occurs in LEs/MVBs. **(A)** Hypothesized model of exosome fusion and cargo release in endosomes. **(B)** Images of R18 dequenching triggered by exosome membrane fusion. R18-dequenching fusion spots (red) were tracked and imaged at the indicated time points via time-lapse microscopy. Scale bars: 5 μm. **(C)** Time-intensity profiles of R18 fluorescence of two representative dequenching spots in experiment (B). **(D)** Membrane fusion signals of exosomes colocalized with the LE marker CFP-RAB7 and the ILV marker CFP-CD63. Dynamic colocalization events of dequenching signals (red) with cellular markers (CFP pseudo-colored green) were tracked via time-lapse microscopy. Scatterplots and Pearson’s correlation coefficients for colocalization are presented below the images. Scale bars: 5 μm. **(E)** Color shift induced by ongoing fusion process of GFP-carrying exosomes prelabeled with self-quenching concentrations of R18 was observed and imaged via time-lapse microscopy. Scale bars: 5 μm. **(F)** Membrane fusion signals of GFP-carrying exosomes colocalized with the LE marker RAB7. Scatterplots and Pearson’s correlation coefficient between the signals of GFP and R18 or the signals of R18 and RAB7 are presented. Fluorescence intensity profiles of GFP, RAB7 and R18 along the indicated white arrow in the ROI are also presented. Scale bars: 10 μm.

We first used time-lapse microscopy to track membrane fusion events in live hepatocytes incubated with macrophage-derived exosomes prelabeled with self-quenching amounts of the hydrophobic dye rhodamine C18 (R18). R18 is commonly used as a fluorescent probe to detect virus-induced membrane fusion. The probe is incorporated into membranes at high concentrations to cause self-quenching, and dequenching of the probe occurs when membrane fusion decreases in density(27, 28). The dequenching signal of membrane fusion was first captured approximately 45 minutes after treating HepG2 cells with R18-labeled exosomes, and fusion events followed within 1 hour (Fig. 5B). The fluorescence intensity profile showed persistent enhanced R18 fluorescence for the fusion spots (Fig. 5C).

EEs and LEs/MVBs are major fusion sites for some viruses to deliver nucleocapsids and release nucleocapsids to the cytosol(46). To locate the exact site at which membrane fusion occurred after exosome internalization, we performed colocalization experiments using a variety of endosomal markers. Endosomal compartments in HepG2 cells were labeled via transient transfection of plasmids encoding CFP-fused markers for EEs (RAB5), LEs/MVBs (RAB7) and intraluminal vesicles (ILVs) in MVBs (CD63). The dequenching signal of membrane fusion was colocalized with the LE marker CFP-RAB7 and the ILV marker CFP-CD63 in live HepG2 cells after treatment with R18-labeled exosomes, while no colocalization was observed with markers for EEs (CFP-RAB5) (Fig. 5D). Hence, LEs/MVBs might be the proper site for the membrane fusion of macrophage-derived exosomes after exosome internalization.

To track exosomal cargo after membrane fusion, the live dynamics of exosomal cargo in hepatocytes were tested by monitoring the membrane fusion events of R18-labeled GFP-carrying exosomes using time-lapse microscopy, with exosomes isolated from GFP-expressing macrophages. At the beginning of the experiment, orange fluorescence was observed at the fusion site due to the combined fluorescence emitted by dequenching R18 inserted into exosome membranes and GFP encapsulated in exosomes. As fusion proceeded, the extreme dilution of the R18-labeled membrane components increased the fluorescence of the gradually exposed GFP. A complete color switch was accomplished when the exosomal cargo GFP was totally uncoated and “released” (Fig. 5E). In addition, confocal images proved again that LEs were the site of membrane fusion for GFP-carrying exosomes (Fig. 5F). The colocalization coefficient of R18 and GFP was approximately 0.9 in HepG2 cells, indicating a high frequency of fusion events among internalized exosomes. Together, these data indicated that LEs/MVBs provided the proper conditions for exosome fusion and cargo uncoating, which might promote exosomal cargo release based on endosome penetration.

### Lysobisphosphatidic acid (LBPA) contributes to exosome fusion and the uncoating of exosomal cargo

Anionic lipids are beneficial for endosome penetration(46). A high concentration of anionic lipids makes LEs a suitable location for endosome leakage via membrane fusion. Notably, the LE-specific anionic lipid LBPA assists as both viruses and delivery vectors to achieve efficient cytosolic access via membrane fusion-induced endosome penetration(43–48).

The accumulation of PKH26-labeled exosomes in the LBPA-rich structure suggested a potential interaction between the two components (Fig. 6A). Partial colocalization between the dequenching R18 of exosomes and LBPA signals indicated the participation of LBPA in the membrane fusion of exosomes in LEs/MVBs (Fig. 6B). To verify the dependence of exosome fusion on LBPA, HepG2 cells were pre-incubated with an anti-LBPA blocking antibody(27, 43), and the dequenching signals of R18-labeled exosomes were tracked via time-lapse microscopy. Pretreatment with an anti-LBPA blocking antibody produced significant inhibition of membrane fusion, as suggested by the decayed R18 dequenching of exosomes (Fig. 6C). The fluorescent intensity profile of tracked fluorescent puncta further manifested the dependence of exosome fusion on LBPA (Fig. 6D). To inquire into the contribution of LBPA to the potential intracellular release of exosomal cargo before lysosomal degradation, we incubated LBPA-blocked HepG2 cells with GFP-carrying exosomes and judged the delivery efficiency of exosomal cargo to lysosomes based on the colocalization efficiency between GFP and lysosomes. The incidence of colocalization increased significantly in cells pretreated with the anti-LBPA blocking antibody, as indicated by the 2-fold increase in the colocalization coefficient (0.5487) in comparison to control cells (Fig. 6E). This finding suggested that some exosomal cargo might escape from endosomes to avoid lysosomal degradation via LBPA-dependent membrane fusion in LEs/MVBs. Unfortunately, the blocking of LBPA in HepG2.2.15 cells led to a four-fold increase in HBV DNA in the supernatant, suggesting that LBPA is closely related to HBV replication (Fig. 6F). This outcome impeded further investigations of LBPA in exosome-mediated antiviral activity transmission. Taken together, LBPA is very important for exosome fusion and the uncoating of exosomal cargo.

**Figure 6.**
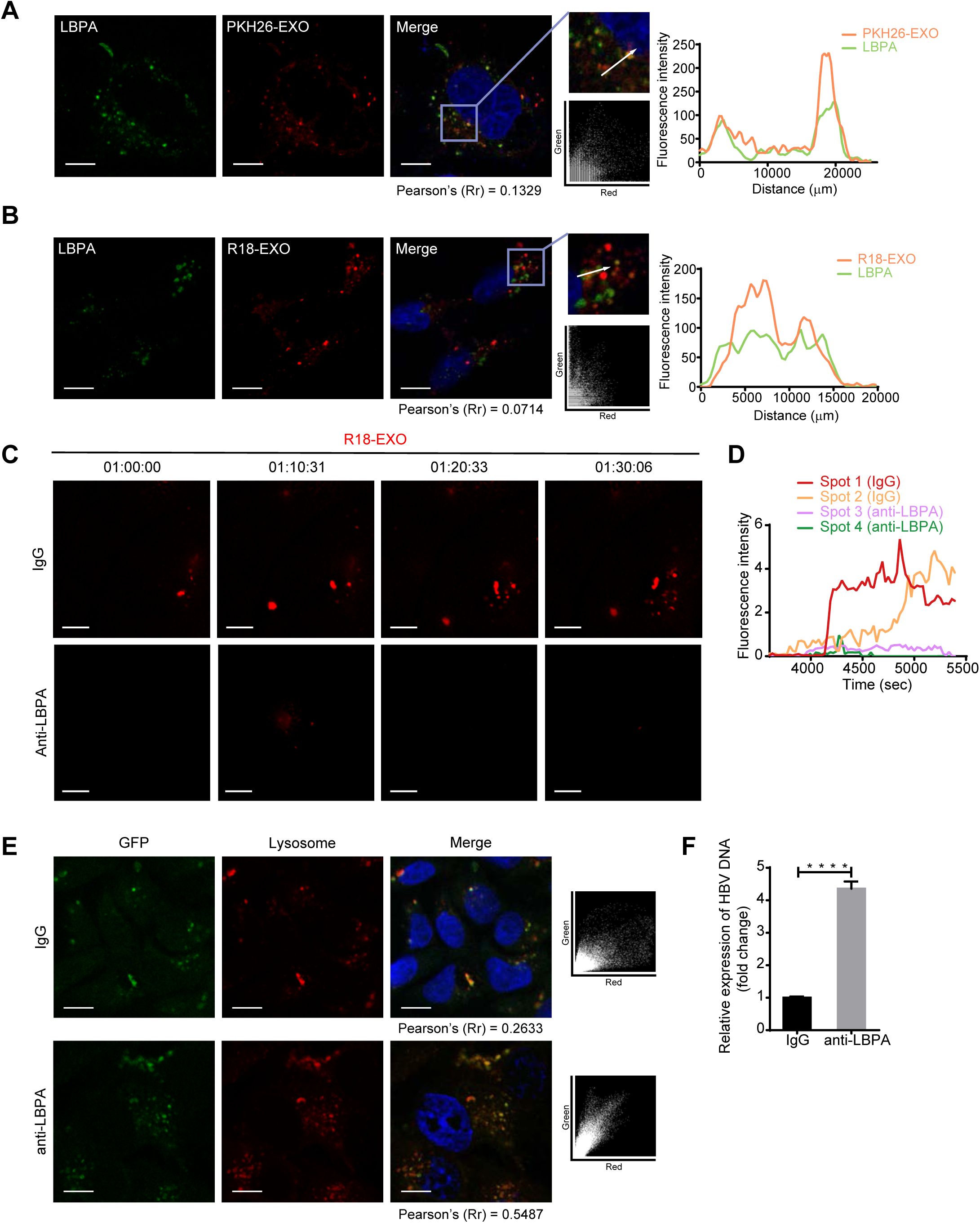
LBPA is required for exosome fusion and cargo uncoating. **(A)** Accumulation of PKH26-labeled exosomes in LBPA-rich vacuoles. Colocalization of PKH26 (red) with LBPA (green) is analyzed as mentioned above. Scale bar: 10 μm. **(B)** Membrane fusion signals of dequenching R18-exosomes colocalized with LBPA. Colocalization of dequenching signals (red) with LBPA (green) is analyzed as described above. Scale bars: 10 μm. **(C)** Inhibition of exosome fusion by antibodies against LBPA. Fusion spots of dequenching R18-exosomes in HepG2 cells pretreated with 50 μg/ml anti-LBPA or anti-IgG overnight were tracked and photographed at the indicated time points. Scale bars: 5 μm. **(D)** Time-intensity profiles of R18 fluorescence of four representative dequenching spots in experiment (C). **(E)**Increase in colocalization of the exosomal cargo GFP with lysosomes after exposure to antibodies against LBPA. HepG2 cells pretreated with anti-LBPA or anti-IgG overnight were incubated with GFP-carrying exosomes in the presence of Lyso Tracker. Colocalization of GFP (green) with lysosomes (red) is analyzed via scatterplots and Pearson’s correlation coefficients. Scale bars: 10 μm.

## Discussion

In this report, we demonstrate that macrophage-derived exosomes utilize virus entry machinery and pathway to proffer IFN-α-induced HBV resistance to hepatocytes. We have presented evidence that macrophage exosomes engage TIM-1, a PtdSer receptor, to enter hepatocytes and undergo rapid CME or sustained macropinocytosis. Our data also suggest that LEs/MVBs are the primary location for LBPA-mediated exosome fusion and accompanying exosomal cargo uncoating for potential intracellular release. The endocytic pathway and membrane fusion in endosomes provide an ideal strategy for exosomes from IFN-α-induced macrophages to deliver antiviral activity and control HBV replication in hepatocytes (Fig. 7).

**Figure 7.**
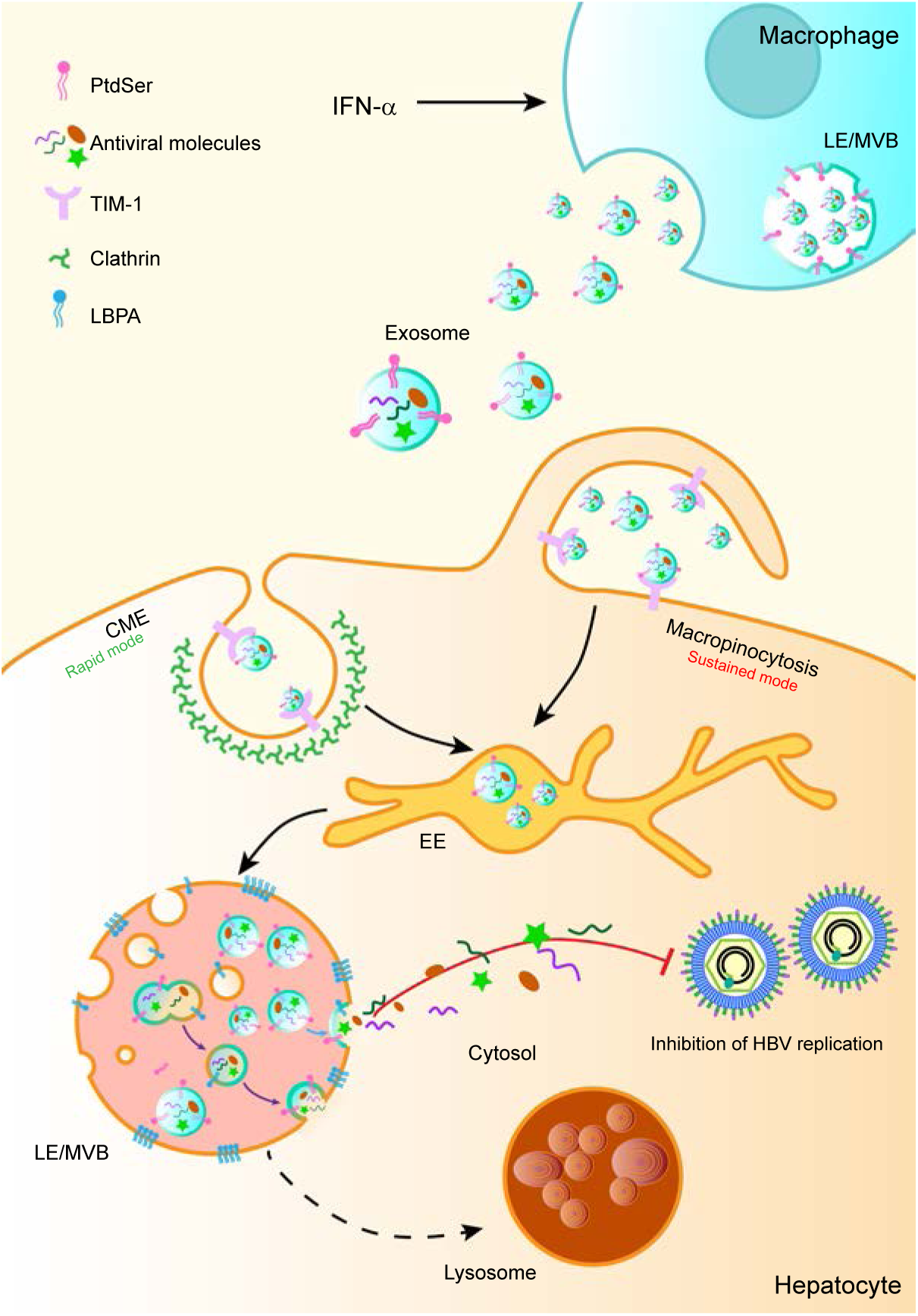
Proposed model of exosome entry and delivery of IFN-α-induced HBV resistance. After binding to TIM-1, exosomes from IFN-α-stimulated macrophages enter HBV-replicating hepatocytes through CME (rapid mode) and macropinocytosis (sustained mode). Endocytosed exosomes traffic to LEs/MVBs and fuse with LBPA-rich ILVs. Trapped antiviral cargo in the ILVs are released to the cytosol via the back-fusion of ILVs with the limiting membrane of LEs/MVBs (violet arrow). Alternatively, ILV-derived exosomes release antiviral cargo via direct fusion with LEs/MVBs (blue arrow).

Exosomes have been shown to interact with membrane receptors on target cells to facilitate subsequent endocytosis(33). Recently, a virus endocytic model — apoptotic mimicry — was suggested to play a role in exosome entry(9, 31). As former ILVs form by inward budding of the LE/MVB-limiting membrane, exosomes are thought to expose PtdSer, an apoptotic marker, on the external leaflet of the membrane and initiate PtdSer receptor-engaged uptake(49). Apoptotic mimicry has been used by hepatotropic hepatitis A virus (HAV) for infection, in which the virus is cloaked in a PtdSer-containing envelope by hijacking the exosome secretion pathway and entering target cells via TIM-1-mediated internalization(31, 50, 51). In this study, we verified PtdSer expression on the external membrane of macrophage exosomes and found that the knockdown of TIM-1 significantly blocked exosome entry and the transfer of IFN-α-induced HBV resistance into hepatocytes. These results indicate that macrophage exosomes may exploit an endocytic strategy similar to apoptotic mimicry as HAV uses to enter cells via TIM-1-mediated internalization. However, the possibility is not excluded that additional receptors may act as co-factors to enhance the attachment of exosomes onto hepatocytes for subsequent entry. Several adhesion molecules enriched on exosome surface, including integrins, immunoglobulins and proteoglycans, are reported to be involved in exosome attachment to cells(9, 21, 33), which implies the necessity of further study on co-receptors.

Adhesion to receptors commonly results in a cellular response of internalization through endocytic pathways(41). Experimental evidence implies important roles for various endocytic pathways in exosome entry, including CME, caveolae-mediated endocytosis, macropinocytosis and phagocytosis(22, 23, 30, 33). It is believed that various combinations of endocytic mechanisms are responsible for exosome entry in different cell types(33). PtdSer exposure is exploited by some viruses as apoptotic disguise which triggers subsequent CME or macropinocytosis for virus entry(31).

According to the results, macrophage exosome entry is sensitive to dynamin and cholesterol inhibitors. Dynamin mediates the fission of endocytic vesicles from the plasma membrane in several endocytic mechanisms, such as CME and caveolae-mediated endocytosis(36). Recent research indicates that dynamin also regulates the closure of circular ruffles during macropinocytosis(37). Cholesterol is an essential constituent of functional domains on the membrane, including lipid rafts and caveolae(40, 41). Cholesterol is required for the formation of endocytic vesicle budding and membrane ruffling(38, 39).

Exosome entry was inhibited by CPZ or CHC knockdown. The rapid accumulation of exosomes and transferrin in the same transport intermediates affirmed that CME plays a role in early exosome uptake by hepatocytes. However, CAV1 knockdown had no effect on exosome internalization. Considering the low expression of CAV1 in HepG2 cells and primary hepatocytes(52), caveolae-mediated endocytosis may contribute little to exosome uptake by hepatocytes.

Furthermore, we found that EIPA, the hallmark inhibitor of macropinocytosis, blocked macrophage exosome entry into hepatocytes. The dependence of exosome entry on PAK1 and PKC was also validated based on the decreased internalization caused by the corresponding kinase inhibitors. In addition, the increasing colocalization of exosomes with dextran during exosome uptake implied that macropinocytosis serves as an efficient alternative route for sustained exosome entry. However, Rac1 and Cdc42, two Rho GTPases that are usually engaged in macropinocytosis, do not appear to be involved in macrophage exosome uptake by hepatocytes. This finding is inconsistent with the interference of exosome entry by EIPA, which inhibits the activation of Rac1 and Cdc42 by altering the sub-membranous pH(42). Therefore, exosome entry into hepatocytes may rely on undefined EIPA-sensitive Rho GTPases. Moreover, Rac1- and Cdc42-independent macropinocytosis is reportedly invoked during influenza A virus (IAV) entry(53). Related studies also showed that circular ruffling and macropinocytosis independent of Rac1 or Cdc42 could be triggered by the non-receptor tyrosine kinase c-src(54). The inhibition of CME or macropinocytosis attenuated exosome-mediated IFN-α-induced anti-HBV transmission, which indicates that exosomes derived from IFN-α-stimulated macrophages utilize both endocytic mechanisms to deliver HBV resistance to HBV-replicating hepatocytes.

Little research to date has focused on the fates of exosomes and exosomal cargo after internalization(9). Endocytosed substances are usually directed to the endosomal system, where they are sorted, processed, recycled, stored and degraded(41). The endosome system is primarily composed of EEs, recycling endosomes (REs), LEs and lysosomes(41). LEs often take the form of MVBs. Invagination and inward budding of the limiting membrane of LEs form ILVs (exosomes) within MVBs(55). Viruses and delivery vectors exploit endosomes for penetration into the cytosol through membrane fusion to deliver viral genomes or biologics(44–46).

Using a live cell imaging system and a fusion probe (R18), we found that LEs/MVBs were also the potential site of exosome fusion initiation, followed by cargo uncoating. Notably, the persistence of R18 dequenching signals for several minutes indicated that exosome fusion was trapped in an endosomal sub-compartment, identical to the colocalization of fusion signals with an ILV marker (CD63) (Fig. 5C and D)(27).

Previous studies have shown that a high concentration of anionic lipids in LEs provides an appropriate environment for endosome penetration(46, 56). It was reported that the presence of anionic lipids in the target membrane promoted membrane fusion efficiency for some enveloped viruses(43, 47, 57). LBPA is a specific anionic lipid in LEs and is thought to promote ILV budding and back-fusion(55, 58) during MVB biogenesis. Research has suggested that the vesicular stomatitis virus (VSV) loads nucleocapsids into ILVs through membrane fusion and penetrates LEs/MVBs through LBPA-dependent back-fusion between the ILV membrane and the endosome-limiting membrane(43). In addition, LBPA is also required for efficient cytosolic access of delivery vectors, including dfTAT and phosphorothioate-modified antisense oligonucleotides (PS-ASO)(44, 45).

Our results showed that the fusion sites of exosomes were colocalized with LBPA. Moreover, LBPA antibodies inhibited the membrane fusion of endocytosed exosomes and accelerated the transport of exosomal cargo to lysosomes. It is possible that some exosomal cargo may avoid lysosomal degradation via LBPA-dependent membrane fusion in LEs/MVBs. Given the above results, we hypothesize that LBPA facilitated the fusion of exosomes from IFN-α-stimulated macrophages with ILVs in LEs/MVBs and that exosomal antiviral cargo are then reloaded into fused ILVs and released after back-fusion with the limiting membrane of LEs/MVBs. As former ILVs formed in LEs/MVBs, endocytosed exosomes with ILV properties may also be qualified for direct fusion with the limiting membrane of LEs/MVBs to release cargo.

Overall, our results illustrate how receptors, endocytic pathways and LBPA-dependent membrane fusion are exploited by macrophage exosomes to deliver IFN-α-induced anti-HBV activities to hepatocytes. This study also highlights the overlap between viruses and exosomes by identifying that the infection strategies of viruses are also applied for exosome entry and exosomal cargo delivery. Dissecting the complete endocytic routes of exosomes may provide a fundamental basis for engineering exosomes as therapeutic vehicles to deliver antiviral molecules with high efficiency.

## Acknowledgments

We thank Zhigang Yi, Xiaonan Zhang for valuable comments and suggestions, and Lu Bai, Xiaoting Du, Yaming Li, Ke Qiao, Shuhui Sun for technical assistance.

This work was supported by research grants from the National Natural Science Foundation of China (81471932), the National Natural Science Foundation (91542207), the National Key Research and Development Program of China (2016YFC1200400) and Shanghai Rising-Star Program (14QA1400700).

The authors declare no competing financial interests.

